# Anomalous diffusion analysis reveals cooperative locomotion of adult parasitic nematodes in sex-mixed groups

**DOI:** 10.1101/2024.04.29.591609

**Authors:** Ruth Leben, Sebastian Rausch, Laura Elomaa, Anja E. Hauser, Marie Weinhart, Sabine C. Fischer, Holger Stark, Susanne Hartmann, Raluca Niesner

## Abstract

Parasitic worms are widespread throughout the world and causing agents of chronic infections in humans and animals. The majority of these pathogens parasitize the gut of the infected hosts, however very little is known about the locomotion of the worms dwelling the gut. We studied the movement of *Heligmosomoides bakeri (*previously *Heligmosomoides polygyrus)*, a natural infection of mice and usually used as animal model to study the roundworm infections. We investigated the locomotion of *H. bakeri* in simplified environments mimicking key physical features of the intestinal lumen, i.e. various medium viscosities, and a periodical intestinal villi topography. The non-periodical nematode motion in these settings could be described by anomalous diffusion theory. Fascinatingly, an oriented, super-diffusive locomotion of nematodes in sex-mixed groups were detected, in the sense of a cooperative, but not collective (swarm-like) migration stimulated by mating and reproduction, while individual nematodes moved randomly, following a diffusive motion regime. High mucus-like medium viscosity and villi topography, representing physical constraints of nematode locomotion, slowed down but did not prevent this cooperative migration. Additionally, the mean displacement rate of nematodes in sex-mixed groups of 6·10^−4^ mm/s in viscous mucus-like medium are in good agreement with estimates of nematode migration velocities between 10^−4^ to 10^−3^ mm/s in the gut. Thus, our data indicate the intestinal nematodes motion to be non-periodic and random but triggered to be oriented by kin of the different sex.

## 1. Introduction

One of the most common parasite infections are infections with parasitic worms also known as helminths. These parasites infect more than ¼ of the world’s population with the roundworm *Ascaris lumbricoides* being one of the most prevalent human helminth infection globally (1). More than 80% of the helminths dwell in the intestine of their hosts, in particular nematodes, the roundworms. The intestine is a very special environment to live in with exceptional physiological and physical characteristics: i) a flow of mucus and ingested food (ingesta) in one direction; ii) elasticity, it can stretch and contract; iii) changes of its surface topology where long villi in the small intestine become shorter towards the large intestine, iv) a viscoelastic mucus layer covering the luminal side, and vi) an environment in which millions of bacteria and possibly other pathogens (protozoa and fungi) coexist with the helminths (2). Here nematodes reproduce and move, however information about their locomotion is scarce.

The locomotion of parasitic nematodes differs from the locomotion of larger organisms, such as fish, and from the locomotion of micro-swimmers such as microscopic parasites (size in the μm range) (3), in terms of the forces involved, the average velocity and the resulting Reynolds number (4). For instance, adult *Heligmosomoides bakeri*, previously named *Heligmosomoides polygyrus* (5, 6), in the following referred as to *H*.*bakeri*, as natural intestinal nematode of mice and model organism for experimental investigation of nematode infections, has a diameter of ≈ 100 μm and a length of several mm to cm (sex-dependent) (7). It typically appears in the coiled posture, whose diameters vary in the range of few 100 μm. The adult worms can be found in the jejunum and distal duodenum of the small intestine during acute infection (14 days p.i.), but during chronicity only be detected in the proximal part of the small intestine (21 days p.i.). Considering typical lengths of murine small intestine segments, we infer that the maximum displacement amounts 10 to 15 cm within up to 7 days (8), resulting into an estimate for the displacement rate of 10^−4^ to 10^−3^ mm/s.

As previously proposed for the free-living nematode *C. elegans*, nematode movement is assumed to be undulatory, being imposed by their vermiform structure with an interior segmented hydroskeleton, surrounded by contractile muscles and inextensible fibers of the exterior cuticle. The forces acting to result into the undulatory motion are alternative muscle contraction and relaxation in the different body segments and the interior pressure of the hydroskeleton, as previously modelled in 2D (9). In addition, a 3D helical motion has been shown for *Nippostrongylus brasiliensis*, however only in viscous media, *in vitro*, with displacement supported by thrust in the viscous medium (10).

Using metabolic imaging based on NAD(P)H fluorescence lifetime imaging of *H. bakeri* in the infected mouse intestine, we could show that the adult worms adapt their bioenergetics to the high energy demand necessary for locomotion against the peristaltic flow, towards the upper part of the intestine (11). Whether *H. bakeri* uses the 2D undulatory or 3D helical regimes to migrate in the intestine, whether this motion follows a periodical pattern, typical for multicellular organisms, and how physical parameters, such as mucus viscosity and intestinal villi geometry influence the *H. bakeri* motion is still unknown. Additionally, the reasons for their migration towards the upper part of the duodenum remain elusive, posing the question whether they compete for scarce resources in the intestine, move randomly, or cooperate.

Here, we isolated adult *H. bakeri* from the small intestine during the 11-28 days p.i. and investigated their locomotion in different artificial environments, to reduce the complexity of the natural habitat and to separately address its individual characteristics, such as the intestinal topography, the medium viscosity or the presence of other individuals of the same species. We found that the nematode locomotion is non-periodical and thus analyzed the nematode trajectories relying on the anomalous diffusion theory, previously used to analyze particle or cell motion (12, 13). We show the anomalous diffusion theory to be applicable for *H. bakeri* motion, as a multicellular organism, and reveal cooperative nematode locomotion in sex-mixed groups.

## 2. Material and methods

### 2.1. *H. bakeri* infection

Wild-type C57BL/6 mice aged 8–10 weeks were purchased from Janvier Labs (Saint-Berthevin, France). All animals were maintained under SPF conditions and were fed standard chow *ad libidum*. Mice were infected with 200 third-stage infective (L3) *H. polygyrus* larvae via oral gavage in 200 μl drinking water. On the dissection days, the mice were sedated by inhalation of isofluorane and sacrificed by cervical dislocation, followed by extraction of the small intestine. Individual female and male *H. polygyrus* worms were isolated from the small intestines of infected mice and were plated out in a 96-well plate containing ∼150 μL/well of RPMI medium. The worms were incubated in serum-free RPMI with 2x PS at 37°C for 24 h to remove remains of mouse tissue, mucus and microbial contaminations prior to imaging experiments. All animal experiments were performed in accordance with the National Animal Protection Guidelines and approved by the German Animal Ethics Committee for the Protection of Animals in Berlin (LAGeSO, G0207/19).

### 2.2. Microscopic and mesoscopic imaging

Microscopic images were taken with the head of a commercial laser scanning microscope (TriMScope II, LaVision Biotec, Miltenyi, Bielefeld, Germany) used in transmitted light, wide field mode. The transmitted light of a white light lamp was collected by an objective (UPlan FLN, 4x, NA 0.13, Olympus, Hamburg, Germany) and detected by a commercial 18.0 megapixel digital reflex camera (EOS600D, Canon, Japan) attached to one of the oculars of the microscope by a specialized adapter (Canon EF-M-Kamerabajonett for LM Digital SLR Adapter, Micro Tech Lab, Graz, Austria) instead of the camera objective. The movies were recorded in time-lapse mode with 25 fps. The image dimensions/ pixel size was determined by the grid spacing (63.5μm) of a square pattern 400-mesh copper grid (3.05 mm diameter, TEM support grid, agar scientific, Stansted, Essex, UK).

Mesoscopic imaging was carried on a standard stereoscope (hund, Wetzlar, Germany) used in transmitted light mode. The transmitted light of a white light lamp was collected by an objective (0.64x or 0.8x magnification) and detected the same way as the microscopic images, by the Canon digital reflex camera attached to the stereoscope ocular with the specialized adapter. The movies were recorded in time-lapse mode with 1 fps. The image dimensions/ pixel size was determined by the diameter (3.05 mm) or the grid spacing (63.5μm) of the square pattern 400-mesh copper grid.

### 2.3. Nematode handling and movement conditions

*H. bakeri* are exothermic organisms, meaning they stop moving when the temperature of the environment is too cold. For imaging, a heating plate with a hole for light transmission was placed under the plates with the *H. bakeri* worms to keep the medium at 31°C.

For the microscopic imaging, the worms were placed individually in a 96-well plate with U-shaped bottom to keep the worms in focus to study the motion patterns as well as to create “constrained conditions” of the movement.

For the mesoscopic imaging, the worms were placed individually or in sex-mixed cohorts of 5-10 animals in a flat-bottomed 24-well plate with (14mm diameter) to create free movement conditions on a “flat plate”, i.e. without constraint during the image acquisition time window.

The worms were kept in RPMI cell culture medium to provide them with nutrients, and when not imaged, they were held in a 37°C incubator. For imaging, the worms were gently placed on the well plates using disposable transfer pipettes with the tip cut off to increase their diameter.

### 2.4. SYOX Green staining

For live/ dead staining, 100μl SYTOX® Green staining (Thermo Fisher Scientific, Darmstadt, Germany) was added in ∼500μl worm suspensions. SYTOX® Green is a green fluorescence nucleic acid stain that does not penetrate living cells and, by that, is an indicator of dead cells or, in our case, small dead organisms. The fluorophore was excited with a mercury vapor lamp, the emission was detected as described above for microscopic imaging after passing a 534/40 nm bandpass filter (AHF analysentechnik AG, Tübingen, Germany).

### 2.5. Alginate mixture

To create mucus-like viscosities, the cell culture medium RPMI was thickened with 2% sodium alginate, following published protocols (10). Sodium alginate is not toxic as it is used in food processing technology and forms a transparent gel in the presence of calcium ions. The resulting viscosity of 2% alginate measured by rotation viscometry (ATAGO VISCO-895, Gimat GmbH Liquid Monitoring, Polling, Germany) is ≃77 mPa·s. This value is similar to that of mucus in the small intestine (14).

### 2.6. 3D printed villi scaffolds to mimic intestinal topography

A porcine small intestine, 28 mm in diameter, was procured from a local meat wholesale market (Rasch, Berlin, Germany) and subjected to decellularization following an established protocol with slight modifications (15, 16). The mucosa and serosa were mechanically removed, and the intestine was subsequently sectioned into smaller pieces and immersed in 1% sodium dodecyl sulphate (Sigma Aldrich) for 2 hours. Following chemical decellularization at room temperature, the tissue pieces were rinsed and incubated with PBS for an additional hour. Subsequently, they were treated with fresh 1% gentamycin/PBS solution for 5 cycles, concluding with immersion in PBS. The resulting decellularized small intestine submucosa (dSIS) was further processed through enzymatic digestion and chemical functionalization for application in 3D printing. For enzymatic digestion, lyophilized dSIS was exposed to acidic pepsin solution (100 mg of pepsin and 1 g of dry dSIS in 100 mL of 0.01 N HCl, pH 2) at room temperature for two days. Pepsin digestion was terminated by adjusting the pH to 9 with 1 N NaOH solution. The digested dSIS was then methacryloylated by sequentially adding methacrylic anhydride (MAAH) six times into the solution over 3 hours (totaling 1 mL of MAAH per 1 g of dSIS), while maintaining the pH at 9 with 1 N NaOH. Subsequently, the pH was adjusted to 7.4, and the resulting dSIS-MA polymer was dialyzed against PBS for one day followed by distilled water for six days before lyophilization. To formulate the dSIS-MA into a bioactive 3D printing resin, it was dissolved in 0.01 N NaOH at a concentration of 1.5 weight-%, along with 1 weight-% of polyethylene glycol dimethacrylate (4 kDa, Sigma Aldrich) and 1 wt% of lithium phenyl-2,4,6-trimethylbenzoylphosphinate photoinitiator (Sigma Aldrich). Intestine-mimicking 3D hydrogel scaffolds (diameter of 11 mm, villi diameter of 300-500 μm and variable height from 0.5 to 2 mm) were designed using Rhinoceros 5 software and fabricated with a visible light DLP printer (Photon D2, Anycubic), employing a 45-second crosslinking time for every 100 μm layer. Following 3D printing, non-crosslinked macromers and dye residues were removed from the samples by excessive immersion in milli-Q water at room temperature.

### 2.7. Image analysis: pre-processing and object tracking

To analyze nematode movement from the acquired time-lapse videos, we first needed to pre-process the data. Using Fiji ImageJ (version 1.54g, (17)), the standard deviation image of the videos were subtracted from the raw data to remove the image background and to retain only the changing/moving parts in the movies. The RGB videos were transformed to 8-bit gray value images and color-inverted for tracking. The full dynamic range of intensity (8-bit) was used for all images, for image standardization. Tracking was performed by TrackMate an ImageJ plug-in (18) and verfied by visual inspection. We used the threshold detector to identify and segment single nematodes in the individual images and a basic Linear Assignment Problem Solver (LAP) for object (nematode) tracking in the videos. The object tracking results were exported and transferred to a custom-written Python Script to create plots of nematode trajectories, all with the starting point at origin (0,0), i.e. rose plots, as well as to calculate the time-averaged mean squared displacements (MSD). Relying on the MSD time-dependence, we analyzed and represented the directionality of nematode movement (see Results section). The exponent of the MSD time-dependence was determined by linear regression (Origin2023, OriginLab, CA).

From each of the tracks, TrackMate determines the track mean speed, the displacement rate and resulting linearity of forward progressions as well as confinement ratio for all conditions. The *track mean speed* is the average of all instantaneous velocities of the track, 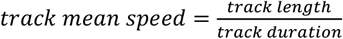. The *mean displacement rate* is the net displacement (air line between start and end of the track), divided by the track duration (see Fig. 3a): 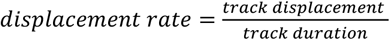. The *linearity of forward progression* is defined as the ratio between the displacement rate and the track mean speed: 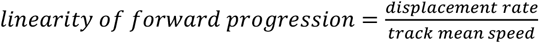. The *confinement ratio* is the ratio of track displacement divided by the track length and indicates how efficient the nematode migrates. It is a unitless measure, taking values between 0 and 1, 0 indicating a fully confined movement, and 1 meaning directed movement along a straight line: 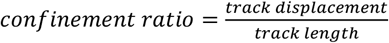. (15)

## 3. Results

### 3.1. The body posture of *H. bakeri* at rest is adapted to the intestinal topography

By comparing the posture of live and dead nematodes in cell-culture medium, we found that a coiled geometry of the worm is the one associated with a minimum of energy requirement, as being the posture of all dead worms (Fig. 1 and Suppl. Video 1). Live nematodes stretch parts of their body to move, and eventually to travel. Labeling with SYTOX Green, a dye which intercalates with the DNA of dead cells, demonstrated the death of the nematodes. The nematodes were killed either by low (−20°C) or high temperature (+100°C), or chemically (4% Neopredisan), to exclude that specific physical or chemical conditions impact on the muscle structure, which defines the body posture.

**Fig. 1:**
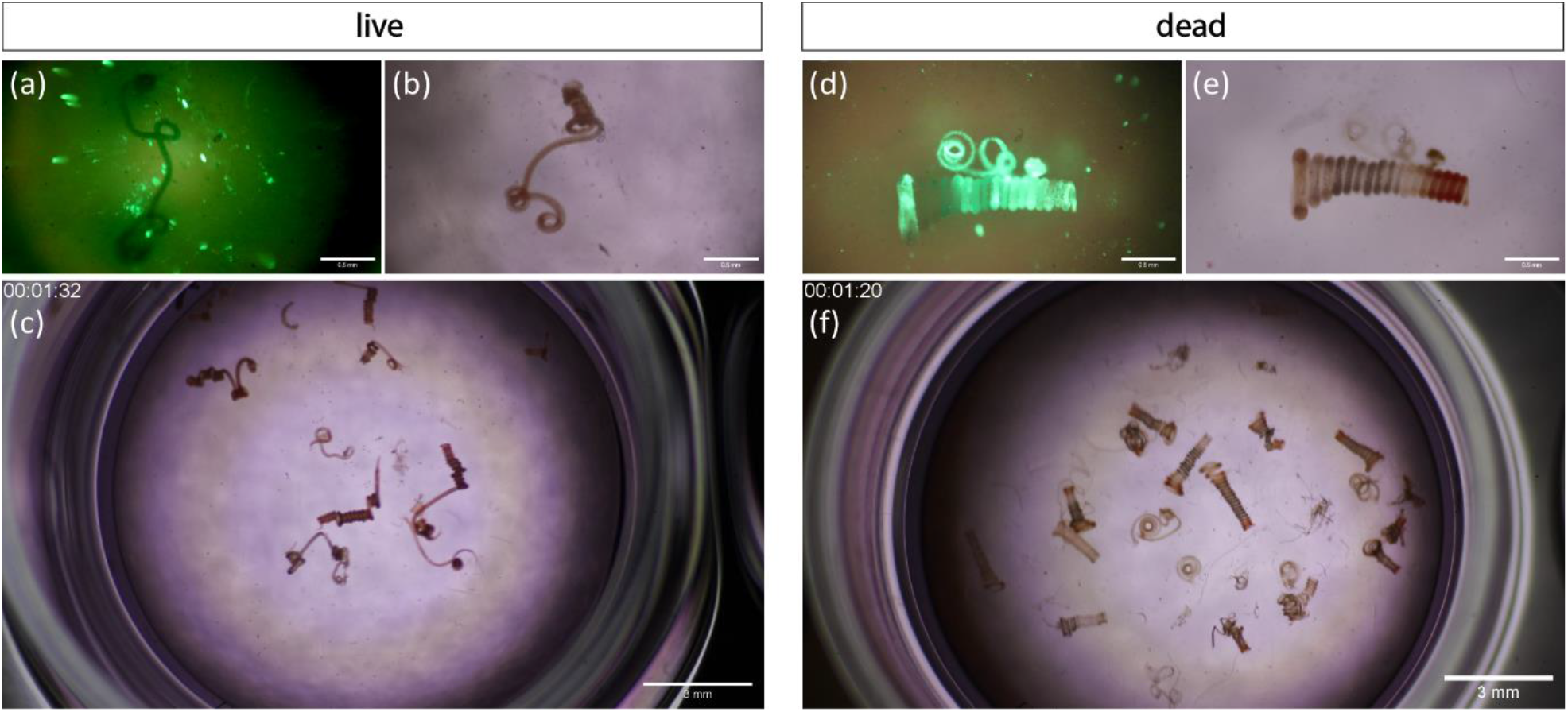
Minimum energy requirement of *H. bakeri* in tightly coiled, helical posture. Comparison of living (left) vs. dead (right) *H. bakeri* worms in cell culture medium heated to 31°C. Top row: Microscopic images at 4x magnification; **(a)** and **(d)** SYTOX® green fluorescence indicates dead cells/organism. **(b)** and **(e)** Transmitted light microscopy of the same individuals a few seconds later than (a)(d). Scale bars indicate 0.5mm. Bottom row **(c)** and **(f)** still images of mesoscopic movies at 0.67x magnification (Suppl. Video 1). Scale bars indicate 3mm. All dead individuals are tightly coiled, in contrast to actively stretching live nematodes.

We found in 111 nematodes, both females and males, that the loops at rest are 295 ± 10 μm (mean value ± standard deviation) in diameter, with the longer females building more loops than the shorter males (7). The loop diameter at rest is slightly smaller than the typical diameter of a villus in the duodenum of mice, determined from two-photon microendoscopic images (300 ± 19 μm, Suppl. Fig. 1).

### 3.2. Non-periodical *H.bakeri* motion can be described by the anomalous diffusion theory

The sequence of still images and the temporal zoom-in from the Suppl. Video 2 (Fig.2) reveal that motion of *H. bakeri* is a series of apparently random stretches and coils that do not result into a closed and periodic motion cycle. Hence, in contrast to other nematodes, such as *C. elegans* or *N. brasiliensis, H. bakeri*, does not move using periodical patterns, typical for multicellular organisms: neither a 2D undulatory nor a 3D helical motion.

**Fig. 2:**
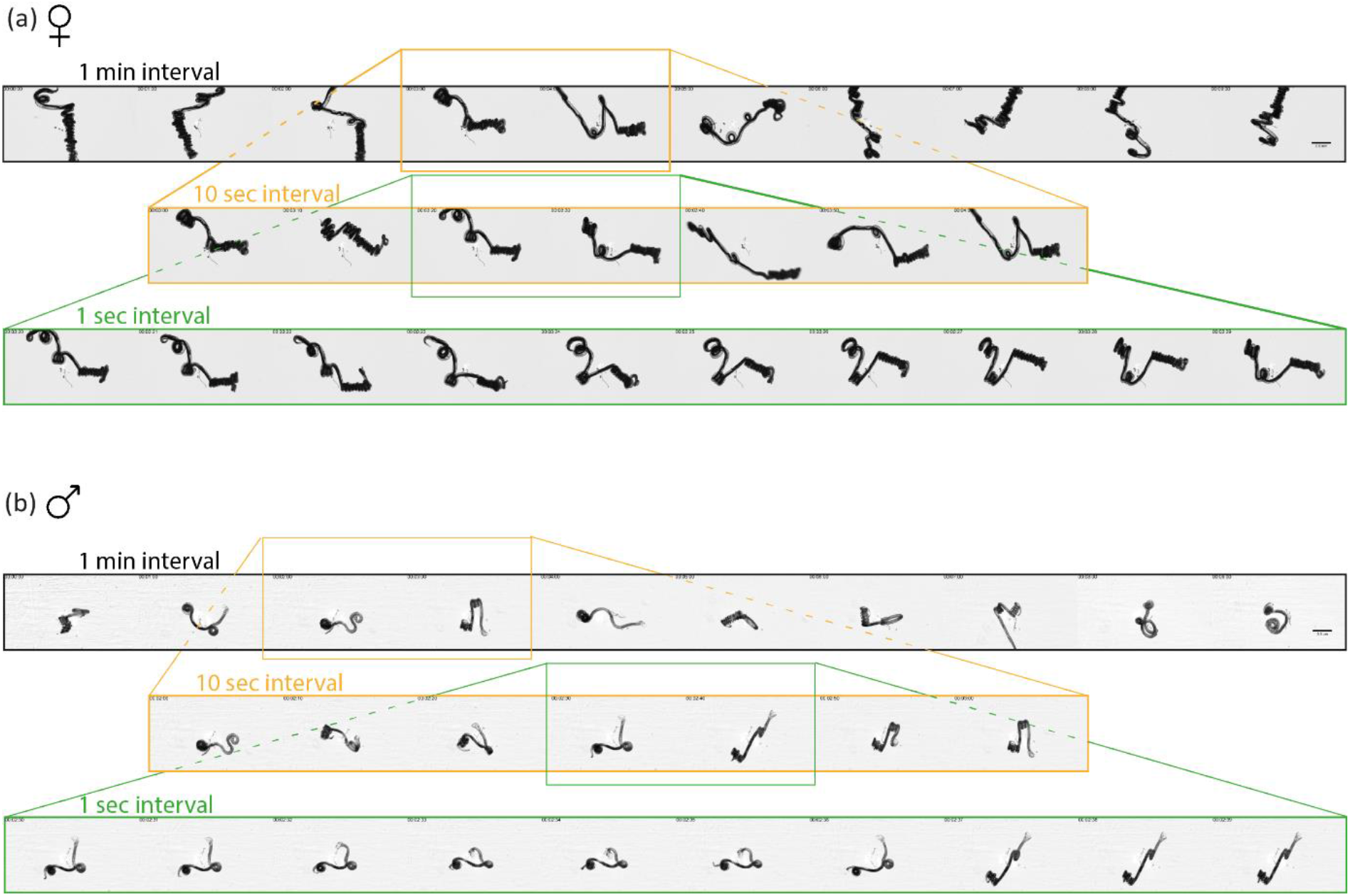
Non-periodic motion pattern. **(a)** and **(b)** Sequences of still images of microscopic movies (Suppl. Video 2) at 4x magnification of adult female **(a)** and male **(b)** *H. bakeri* worms in cell culture medium heated to 31°C. Sequences have different temporal intervals: Top row (black frame) – 10 min sequence with 1min interval time; Middle row (orange frame) – 1min zoom-in sequence with 10sec interval time; Bottom row (green frame) – 10 s zoom-in sequence with 1 s interval time (1 Hz). Time shown is mm:ss. Scale bars indicate 0.5mm and apply for all shown sequences. The movement is irregular, meaning no motion pattern is repeated as e.g. in a sinusoidal (2D) or helical (3D) movement.

As no clear periodical pattern of neither female nor adult male worms could be identified, we hypothesize that the motion of *H. bakeri* can be described by anomalous diffusion.

The diffusion theory describes stochastic molecular motion (Brownian motion), following Fick’s laws. It has been extended with the concept of anomalous diffusion, to describe the motion of molecules, but also of cells in tissues (12, 13) or of unicellular organisms (19, 20), in the presence of additional conditions, such as confinement or other types of transport processes. The time-averaged mean squared displacement (MSD) is a measure of the distance that a object travels on average in the lag time τ = *k* · Δ*t* (21) and is defined as:

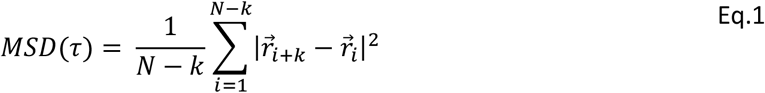

Where 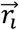 (with *i* = 1,…, *N* − *k* denoting the successive frames of a track and with index *k* = 1,…, *N* − 1) is the position of the object (worm) at the i-th time point,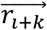 is the position of the object at the (i+k)-th time point, delayed by τ = *k* · Δ*t*, and Δ*t* the constant time span between two succesive frames (Fig 3a).

**Fig. 3:**
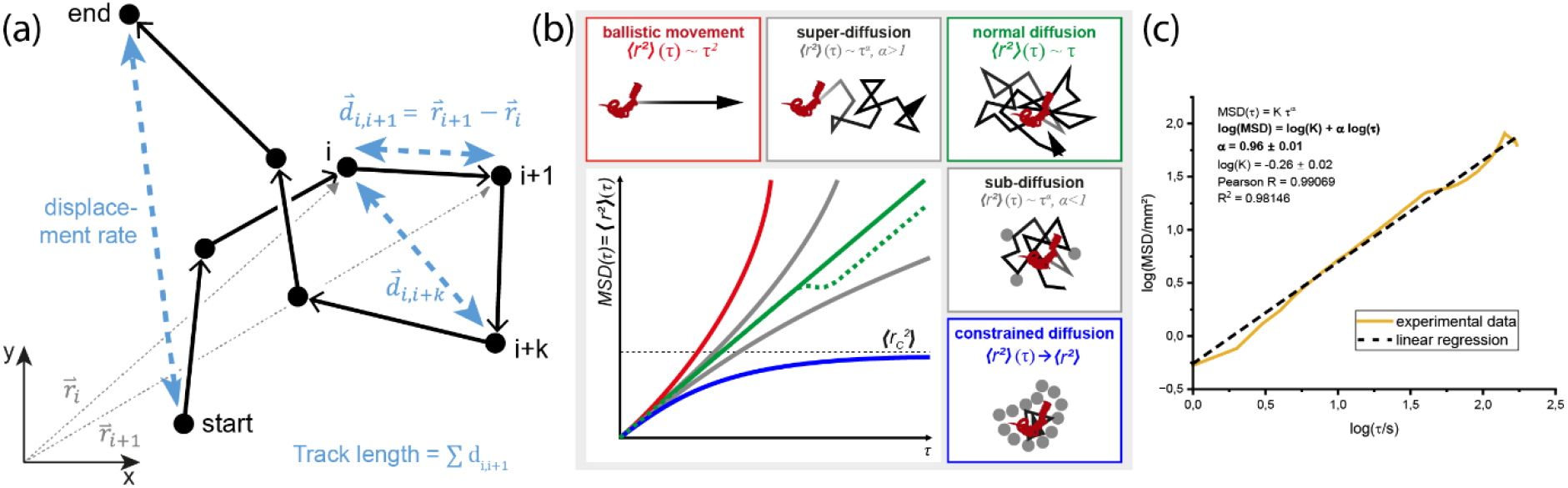
Anomalous diffusion theory: motion regimes determined by the time-dependence of the time-averaged mean square displacement MSD(τ). **(a)** Sketch of relevant vectors and distances to determine MSD(τ). These also form the basis of the different speed measures as well as confinement ratio and linearity of forward progressions. Track displacement is the “air line” from starting point to end point of the track. Total distance traveled is the sum of all distances d_i,i+1_ between time points i and i+1, r_i_ is the position at time i. **(b)** Movement categories according to the diffusion theory: clockwise from upper left to bottom right: ballistic movement (α=2, red); super-diffusion (α>1); normal diffusion (α=1, green, continuous line: one diffusive component, dashed line: two diffusiv components τ_1_ + τ_2_); sub-diffusion (α<1); motion under constrained condition (α=0, blue). Graphic adapted from (22). **(c)** Representative linear fit (black, dashed line) of an experimental MSD time-dependence (orange curve) in double-logarithmic representation, used to determine the slope α, i.e. the exponent in non-logarithmic representation Eq. 2.

Following the anomalous diffusion theory, the time-dependence of the time-averaged mean squared displacement MSD of an object is generally described by:

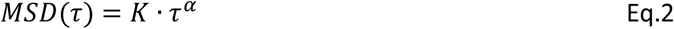

with *K* being a constant with different meanings for the different motion regimes, τ being the lag time and α being the exponent, which describes the motion regimes. The motion regimes (Fig.3) described by α are:

i. confinement/ constrained conditions, i.e. motion but no displacement: α = 0, for which MSD approaches a constant over time,
ii. sub-diffusive regime, i.e. random motion with obstacles: 0 < α < 1,
iii. diffusive motion, stochastic motion following Fick’s laws, i.e. Brownian motion: α = 1,
iv. super-diffusive regime, meaning an oriented motion, as an overlapping directed and random motion: 1 < α < 2,
v. ballistic motion, i.e. straight directed locomotion: α = 2.

First, we verified whether the anomalous diffusion theory also applies to *H.bakeri* nematode motion, i.e. large multi-cellular organisms. Nematode motion in U-shaped dishes serves as a proof-of-principle for confinement, in contrast to random free nematode motion on flat dishes, both in aqueous medium (Suppl. Video 2 and Suppl. Video 3).

By comparing the time-dependence of MSD of both single male and female nematodes under partially confined conditions, i.e. when placed on a U-shaped well in aqueous media, and to their free movement on a flat well, much larger than their size, we found that the theory of anomalous diffusion reliably describes the nematode motion (Fig.4). The exponent α of the linear fit of the time-dependent MSD for partially confined nematode motion is 0.16±0.11 (n = 10 worms, i.e. 5 males, 5 females), with no difference between males and females (Fig. 4a, d, f). Moreover, the corresponding rose plots (inlay in Fig 4d) shows no displacement for all individuals. In contrast, for freely moving nematodes on a flat plate the exponent α increases significantly to 1.01±0.14 (n = 10 worms, i.e. 5 males and 5 females), indicating a diffusive motion regime (Fig. 4b, c, e, g), resembling random motion, similar to the stochastic Brownian motion of molecules at thermic equilibrium. However, sex-dependent analysis of the MSD reveals a shift towards the sub-diffusive regime for females (α = 0.80±0.09) and the super-diffusive regime for males (α = 1.22±0.11), possibly due to limited number of investigated nematodes. Additionally, rose plots of freely moving nematode trajectories show no preferential orientation of locomotion, i.e. displacement (Fig. 4g).

**Fig. 4:**
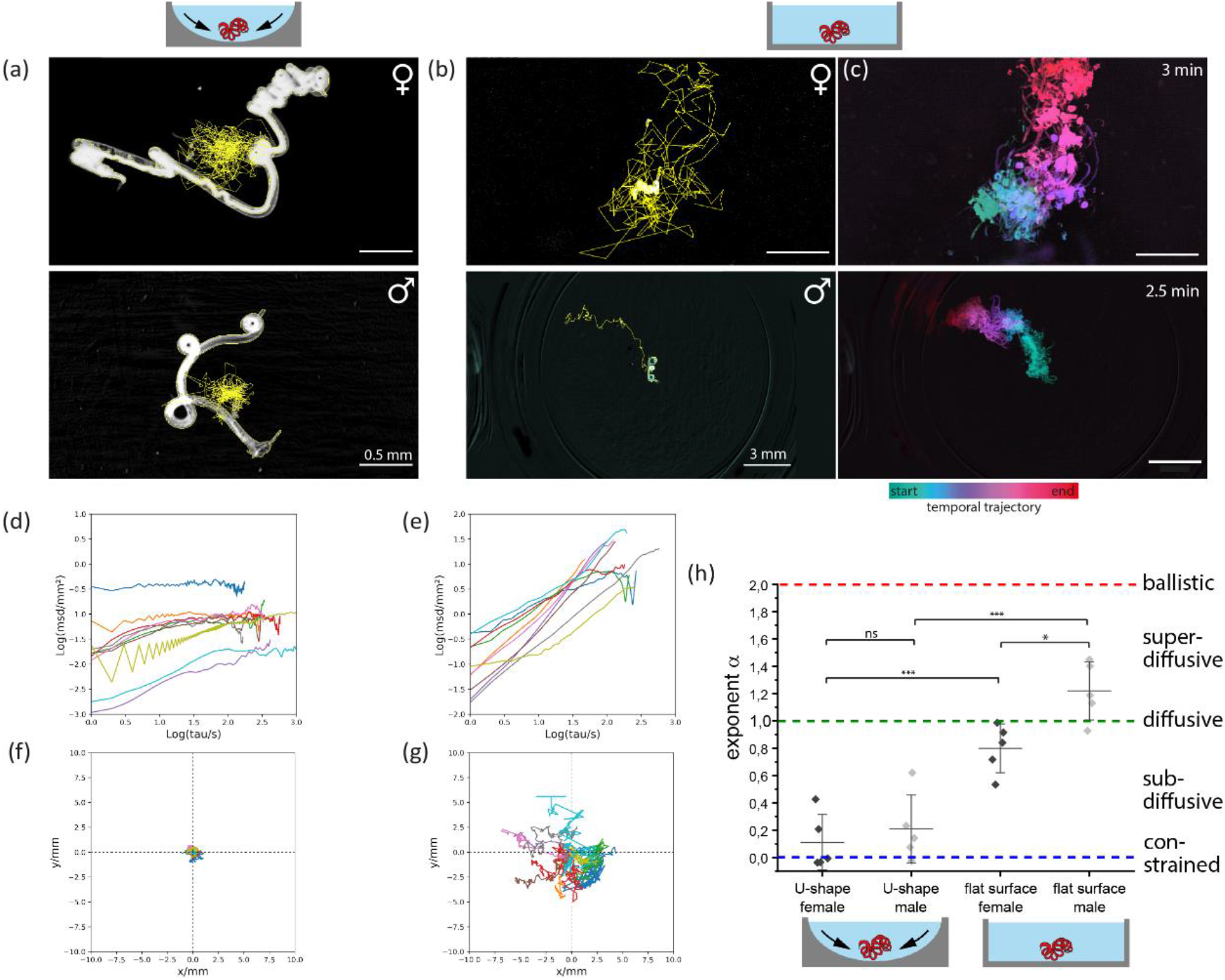
Analysis of nematode motion regimes under confined and free-movement conditions. Single adult *H.bakeri* worms in aqueous cell culture medium heated to 31°C; top row shows exemplary images of a female worm, bottom row of a male worm (one of five per sex and condition). **(a)** Representative microscopic image of one worm (of five) each sex under constrained conditions in a dish with u-shaped bottom with 4x magnification, scale bars are 0.5 mm. **(b)** Motion of one worm (of five) each sex on a flat bottom plate; 4x magnification (Suppl. Video 3), scale bars indicate 3 mm. Yellow lines in (a) and (b) show nematode trajectories (tracking analysis by TrackMate). **(c)** Color-coded temporal trajectories of the data shown in (b). **(d) and (e)** Time-averaged mean square displacement (MSD) of all tracks: **(d)** corresponds to motion under confined conditions, **(e)** and to random, free motion on a flat plate. **(f) and (g)** To (d) and (e) corresponding rose plots, i.e. plots of nematode trajectories, with the starting point set to the origin (0,0). **(h)** Exponent α resulting from the linear fit of the double-logarithmic time-dependent MSD, displayed per sex and condition. α indicates the motion regime, according to the anomalous diffusion theory. Statistical significance is represented by p≤0.001: ***; p≤0.05: *; p>0.05: ns, using the one-way ANOVA with Bonferroni’s post-hoc test. Under unconfined conditions the non-periodic motion of *H. bakeri* results in a forward movement.

### 3.3. Super-diffusive regime of *H. bakeri* motion in sex-mixed cohorts hints towards cooperative locomotion, despite physical intestinal constraints

In order to investigate the interaction between nematodes of different sexes and the impact on their locomotion, we compared the motion of sex-mixed cohorts of nematodes on the flat plate, in aqueous medium (Fig.5a,d,g,j), (each group with n = 5 to 6 females and n = 5 to 6 males) to the motion of single nematodes, under the same conditions. We found an increase of the exponent α from 1.01±0.14 (diffusive regime) to 1.26±0.15 (super-diffusive), for n = 21 worms (Fig.5m). Additionally, all nematodes migrated to the same area 4 minutes after being randomly placed on the well surface (Suppl. Video 4). These observations hint towards a cooperative locomotion of the worms. When characterizing the anomalous diffusion motion regimes in a sex-dependent manner, we found that the sub-diffusive motion regime of single females (α=0.80±0.08, n = 5) turns into a super-diffusive regime of females in sex-mixed groups (α=1.22±0.17, n = 11). In contrast, the super-diffusive motion regime of single males (α=1.22±0.11, n = 5) is preserved also in sex-mixed groups (α=1.29±0.12, n = 10). This may indicate that females rather than males concur to the overall cooperative motion.

The natural habitat of *H.bakeri* is the small intestine, which luminal-sided surface has a villi structure, that are protrusions of the *lamina propria mucosae*. These villi increase the surface area of the intestine for efficient nutrient absorption form the ingesta. We mimicked it’s geometry with 3D printed scaffolds of extracellular matrix-based material (villus diameter 300 μm, gradient in the height 0.5 to 2.0 mm) to investigate the influence of the topology on the worm movement. We found that this topography impedes the nematode motion in sex-mixed groups (Fig.5b,e,h,k), independent of villus length, as we detected a sub-diffusive regime of nematode motion in sex-mixed cohorts when placed on the 3D scaffolds in aqueous media, with α = 0.63±0.13 (n = 10 worms, 5 males and 5 females, Fig. 5n), within 15 minutes observation time.

**Fig. 5:**
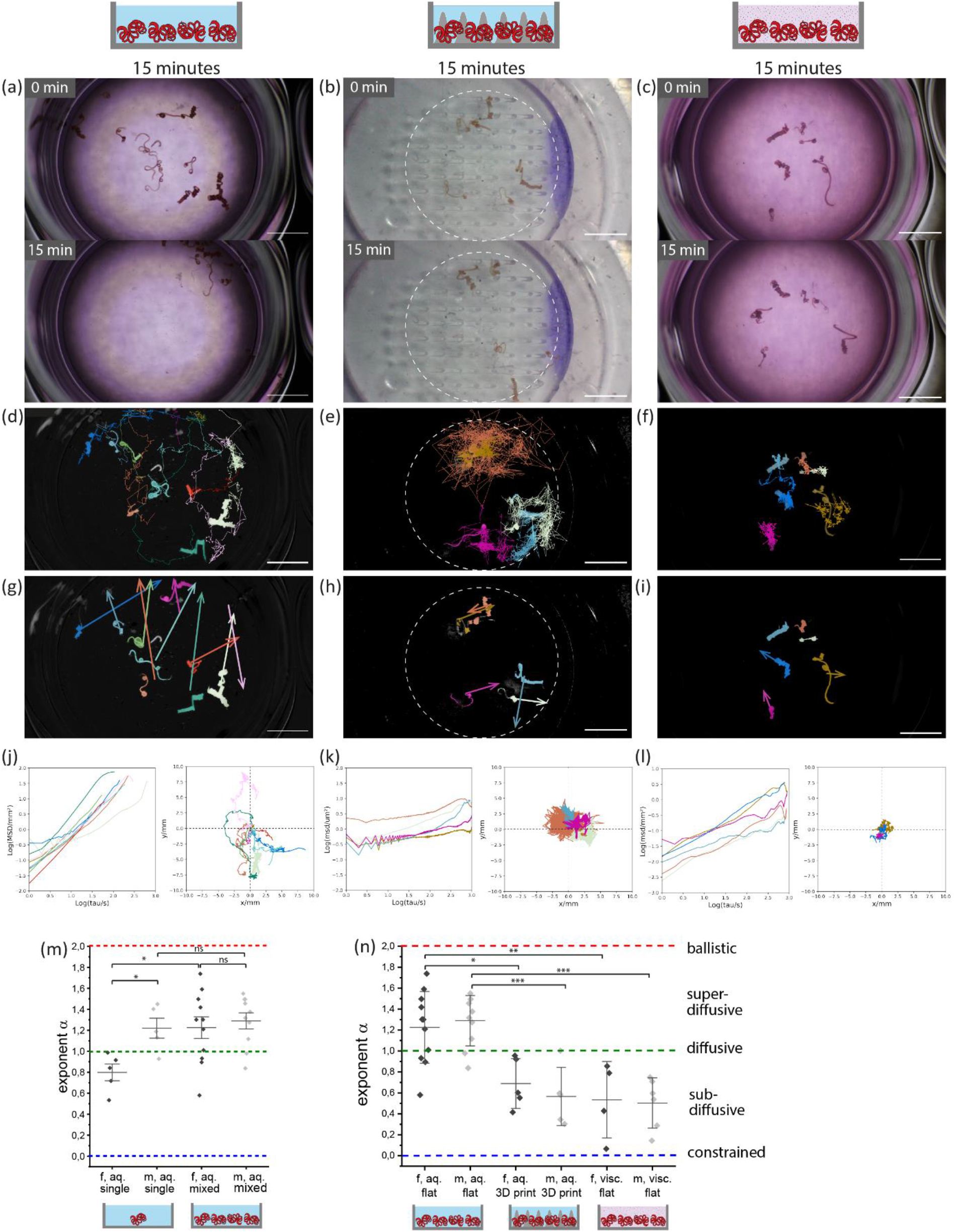
Super-diffusive, oriented motion of *H. bakeri* in sex-mixed cohorts indicates cooperative locomotion, being subject to physical constraints in the intestine. Analysis of the movement of adult *H. bakeri* worms of both sexes in cohorts and in different environments, at 31°C, observed over 15 min: **(left - a**,**d**,**g**,**j)** in aqueous cell culture medium on a flat plate; **(middle – b**,**e**,**h**,**k)** in aqueous culture medium on 3D printed villi structure, dashed circle indicates the area with printed villi; **(right – c**,**f**,**i**,**l)** in viscous cell culture medium thickened with 2% alginate on a flat plate. **(a-c)** Still images of mesoscopic movie at 0.67x magnification (Suppl. Video 4). Scale bars indicate 3mm. **(d-f)** Still images of processed data shown in (a-c) with segmented and tracked worms (TrackMate), entire trajectories are shown. **(g-i)** Still images of processed data shown in (a-c) with the displacement vectors of segmented and tracked worms from (d-f). **(j-l)** Time-averaged mean square displacement (MSD) of the tracks (left) and corresponding rose plots (right). **(m)** and **(n)** Exponent α of the fit of the MSDs per sex under the above presented conditions: **(m)** Single male and female worms vs. female and male worms in a cohort on a flat plate in aqueous medium. **(n)** Female and male worms in a cohort: left - on a flat plate in aqueous medium, middle - in aqueous culture medium on 3D printes villi structure, right - in cell culture medium thickened with 2% alginate on a flat plate. Statistical significance is represented by p≤0.001: ***; p≤ 0.01: **; p≤0.05: *; p>0.05: ns, using one-way ANOVA with Bonferroni’s post-hoc test.

The *lamina propria mucosae* is covered with a layer of viscous-elastic mucus, the viscosity of which is 3mPa·s < η < 100mPa·s (depending of the layer depth (14)) and, thus, higher than aqueous medium (η≃ 1mPa s). We imitate the viscosity of intestinal mucus by 2% alginate in the aquesous medium (η≃77 mPa s, measured by rotation viscometry). We analyzed the motion of sex-mixed cohorts on a flat well, in this mucus-like medium and found that it also impedes the nematode motion (Fig.5c,f,i,l). Again, it resembles sub-diffusive features (α = 0.52±0.14, n = 10 worms, 5 males and 5 females) within 15 minutes observation time (Fig.5n).

For both constraining conditions, the villi-like scaffolds and in mucus-like viscous medium, we found no significant differences between male and females motion regime in sex-mixed cohorts (Fig.5n).

Notably, we found that the worms in sex-mixed cohorts in viscous medium (2% alginate) do migrate to the same area, when we observed them over 45 minutes (Fig.6). However, their motion in viscous medium is slower than in aqueous medium, with a mean displacement rate of 0.0006±0.0004 mm/s compared to 0.03±0.02 mm/s. Additionally, we found an increase of exponent α to 0.96±0.14 (n = 7 worms, 4 males, 3 females), when analyzing the time-dependence of MSD over 45 minutes (at 1 Hz image acquisition rate) instead of 15 minutes (at 0.1 Hz acquisition rate). Thus, the nematode motion regime is dependent on the timescale, which was not expected from the anomalous diffusion theory. Altogether, these results indicate that nematodes can overcome physical intestinal-like constraints to migrate in a cooperative manner.

**Fig. 6:**
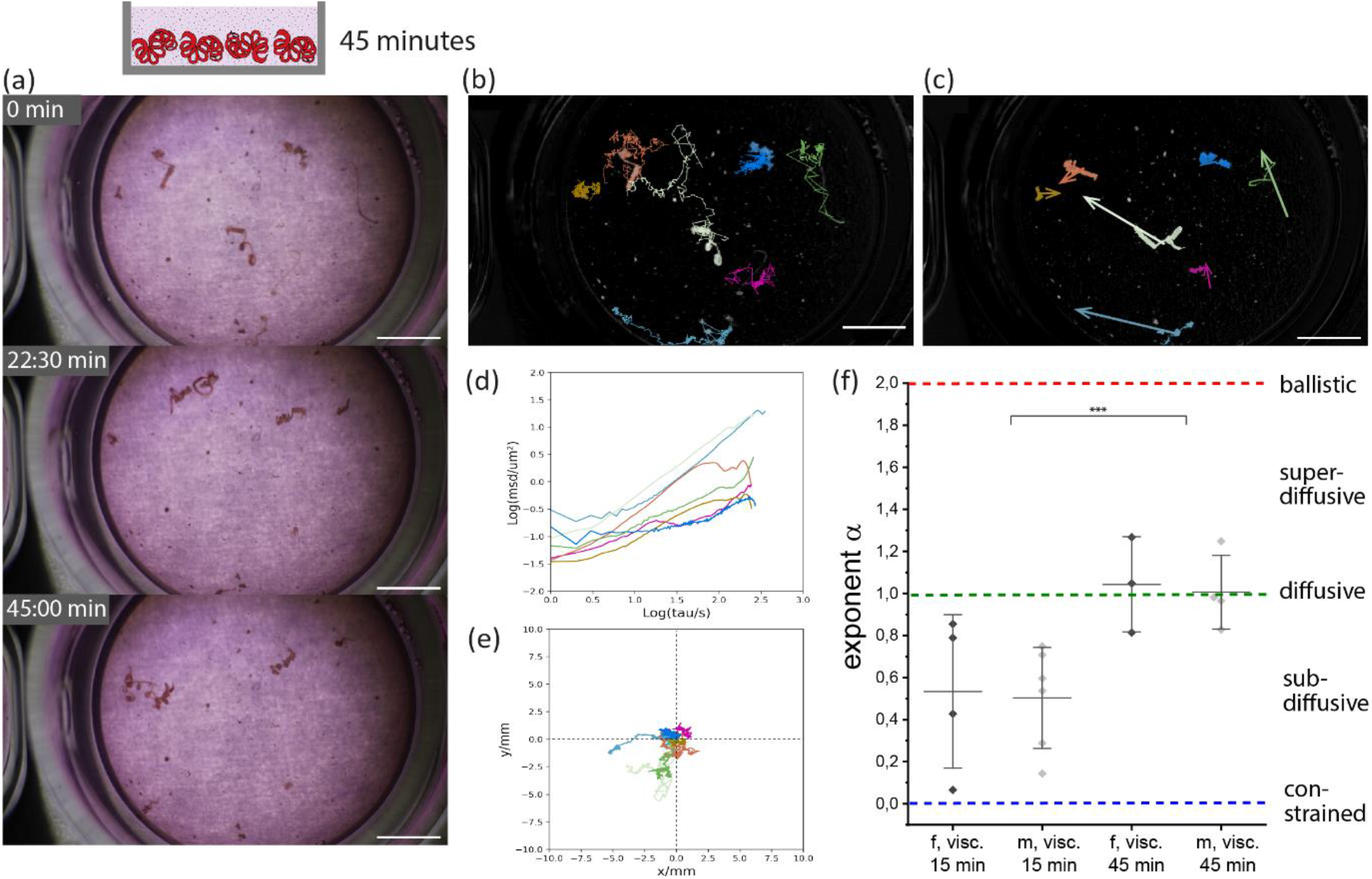
Increasing orientation of nematode motion on larger timescales, despite constraints in mucus-like environment. Analysis of the movement of adult *H.bakeri* worms of both sexes in a cohort in viscous cell culture medium thickened with 2% alginate on a flat plate heated to 31°C. **(a)** Still images of 45min mesoscopic movie at 0.67x magnification (Suppl. Video 5). **(b)** Still image of processed data shown in (a) with segmented and tracked worms (by TrackMate), entire tracks are presented. **(c)** Still image of processed data shown in (a), with displacement vectors of segmented and tracked worms (b). Scale bars in (a), (b) and (c) indicate 3mm. **(d)** Time-averaged mean squared displacement (MSD) of the tracks and **(e)** corresponding rose plots. **(f)** Exponent α of the fit of the MSDs per sex under the above presented conditions: left - within 15min; right – within 45 min. Statistical significance is calculated for the entire cohorts (not separated according to sex) and represented by p≤ 0.01: **, Student’s T-test. Individual worms are able to overcome the constraints by viscosity within 45 minutes and migrate towards each other.

To substantiate the results obtained by MSD and presented above, we finally compare the track mean speed, i.e. the average instantaneous speed of a track, the mean displacement rate and resulting linearity of forward progressions as well as confinement ratio for all conditions (Table1).

As shown in Table 1, we found a high variability of nematode motion under the same experimental conditions. However, the mean displacement rate values show that the worms are several orders of magnitude quicker, when moving freely, i.e. in aqueous medium, on flat plate, as compared to physical constraints, i.e. when placed on the 3D printed villi scaffolds or in mucus-like viscous medium. In contrast, the track mean speed stays the same within the error margins for worms moving in aqueous medium, both on the flat plate and on printed villi scaffolds, but it is decreased in viscous medium. The confinement ratio and linearity of forward progression values confirmed the constrained nematode movement of 3D printed villi scaffolds and in viscous medium.

**Table 1:**
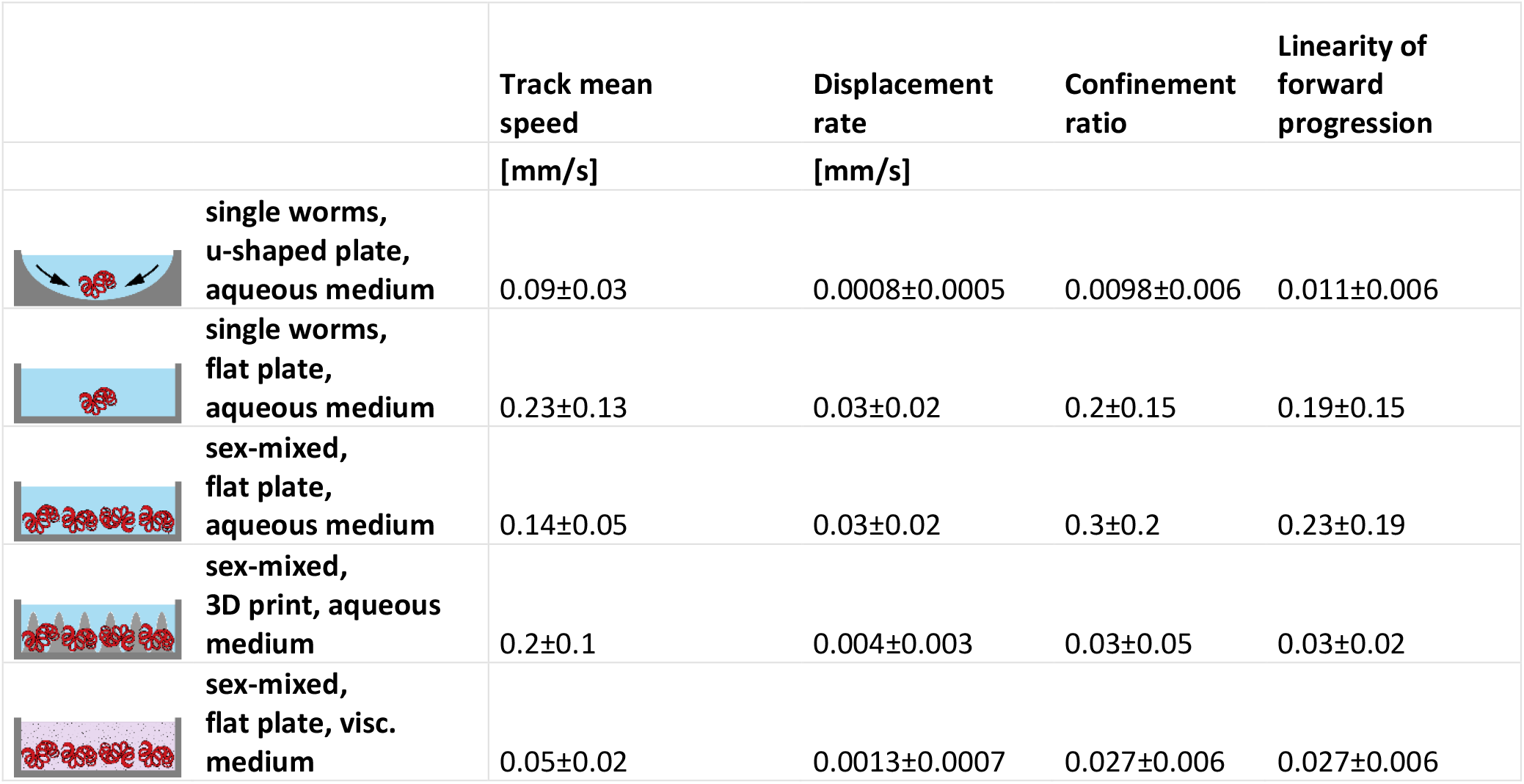
Track mean speeds, displacement rate, confinement ratios and linearity of forward progressions for all conditions.

|  | Track mean speed | Displacement rate | Confinement ratio | Linearity of forward progression |
| --- | --- | --- | --- | --- |
|  | [mm/s] | [mm/s] |  |  |
| single worms, u-shaped plate, aqueous medium | 0.09±0.03 | 0.0008±0.0005 | 0.0098±0.006 | 0.011±0.006 |
| single worms, flat plate, aqueous medium | 0.23±0.13 | 0.03±0.02 | 0.2±0.15 | 0.19±0.15 |
| sex-mixed, flat plate, aqueous medium | 0.14±0.05 | 0.03±0.02 | 0.3±0.2 | 0.23±0.19 |
| sex-mixed, 3D print, aqueous medium | 0.2±0.1 | 0.004±0.003 | 0.03±0.05 | 0.03±0.02 |
| sex-mixed, flat plate, visc. medium | 0.05±0.02 | 0.0013±0.0007 | 0.027±0.006 | 0.027±0.006 |

## 4. Discussion

Gastrointestinal nematodes account for the vast majority (about 80%) of parasitic worm infections, affecting approximately a quarter of the world’s population, in particular in tropical and sub-tropical areas as well as farm and wild animals all over the world. These infections are of considerable global concern leading to millions of disability-adjusted life years (DALY) of infected humans and to huge economic losses in animal husbandry. *Heligmosomoides bakeri*, naturally infecting mice, represents an easily accessible mouse model to study parasitic nematode infections. Briefly, after oral uptake, the larvae invade the small intestinal submucosa before re-emerging and dwelling the gut lumen as long-lived adult worms, leading to chronic infections. Previous studies mostly focused on the persistence mechanisms of parasitic nematodes and their efficient immunomodulation resulting in chronic worm persistence in immunocompetent hosts. This included the immunoregulation of innate host immune cells (23-25), as well as of adaptive immune responses (26-29) and the significant regulation of unrelated autoimmune responses (30-33). Moreover, nematodes induce epithelial immunological quiescence at transcriptional level (34), alter the composition of the microbiota and release antimicrobial factors (2, 35, 36), even promoting the production of antimicrobials by host cells, potentially thwarting efficient anti-parasite immune responses (35). However, despite the close coexistence of intestinal nematodes with millions of bacteria in a mucus-rich environment, induced by the infection itself (2, 37), little or nothing is known about the locomotion of adult worms in the gut, allowing them not only to persist but also to mate and reproduce in the specific host niche, i.e. the small intestine. In particular, the impact of physical conditions such as mucus viscosity and luminal topology and of nematode co-existence, i.e. social condition, on their locomotion is not clear.

By analyzing the posture (geometry) of nematodes as compared to the intestinal topography, consisting of periodically organized villi, we found that the coiled body of nematodes at rest, with no energetic needs, is adapted to the villus geometry in the duodenum. We suggest that this adaptation ensures nematode attachment to the villi and, by that, resistance against the peristaltic flow in the host intestine. The coiled geometry continuously changes in live, motile nematodes, however, not following any obvious periodical pattern, as expected from other nematodes (*C. elegans* (38, 39) or *N. brasiliensis* (10)). Relying on the anatomical fact that longitudinal muscles surround the tubular body of the nematodes directly under the cuticula, we infer that these muscles have a helical geometry at rest, which leads to the coiled body posture, and that the worms need to actively stretch against this internal tension. Some of these muscles contract locally, inducing stretching of the neighboring, connected muscles. This differs from *C. elegans*, for which alternate contraction of the muscles on transversally opposite sides of the body, lead to a 2D undulatory motion.

To be able to determine the effects of physical cues, i.e. medium viscosity and environment topography, and of social cues, in sex-mixed groups, on nematode locomotion, we investigated nematode motion in environments with simple geometry, in which these cues could be individually controlled. The observed motion does not resemble the complexity of nematode movement in the mouse intestine but provides us with insight into its specific aspects.

By analyzing the time-average mean squared displacement of nematode trajectories moving under confined conditions as compared to those freely moving on a flat surface, we found that the anomalous diffusion theory applies to their non-periodical movement. Relying on this theory, we could show that freely moving nematodes in sex-mixed groups can be described by a super-diffusive regime, in contrast to single nematodes, with differences between sexes. As we found that randomly distributed nematodes on a flat surface migrate to the same spot, we infer that the super-diffusive motion is of cooperative, social nature, but not collective (swarm-like) as their motions are not synchronized. This is possibly triggered by mating and reproduction, similar to the effect of pheromones in *N. brasiliensis* (40). Notably, we observed a similar cooperative motion also for other nematodes, i.e. the human and porcine intestinal nematode *Ascaris suum* (unpublished data), indicating that this may represent an overarching nematode adaptation.

Moreover, we found that higher medium viscosity, similar to that of mucus, and intestinal-like villi topology, lead to sub-diffusive nematode motion in sex-mixed groups, suggesting that these physical cues represent motion constraints. However, these physical constraints can be overcome, as we found an increase in orientation of locomotion when extending the observation time-window of nematodes in mucus-like viscous medium.

As we observed that nematodes attempt to coil around the 3D printed villi, sporadically attaching to those for short periods of time (Suppl. Video 6), the sub-diffusive nematode motion may be also explained by short time spans, during which the nematodes pause and inspect the villus structure. In contrast to intestinal villi, on the acellularized 3D printed villi structure we did not long-term nematode attachment. Possibly, this is due to insufficient adhesion of the nematodes on the smooth surface of the 3D-printed villi compared to that of natural cellularized intestinal villi, as microvilli and glycocalyx on epithelial cells lining the villus create a rough surface at nm scale (41). The presence of epithelial cells on the villi may be also required for long-term nematode attachment as the parasite feed on these host cells and not on other nutrient sources such as ingesta or blood (42).

Notably, the migratory velocity, i.e. displacement rate (6·10^−4^ mm/s), of nematodes in viscous mucus-like medium is in good agreement with the migratory velocity of 10^−4^ to 10^−3^ mm/s we estimated for worms travelling maximum several tenths of cm through the intestine, over a time span of several days, starting from the time point the adult worms emerge from the submucosa into the lumen and ending at the time point when they arrived at the upper part of the duodenum.

Concluding, by analyzing nematode motion relying on the anomalous diffusion theory, we found that nematodes migrate in a cooperative manner, in sex-mixed groups, suggesting adaptation triggered by mating and reproduction, with high mucus viscosity and intestinal villi topology, slowing down, but not preventing the nematodes from gathering.

## Author contributions

Conceptualization: RL, SH, SCF, HS, RN; Investigation: RL; Data curation: RL, RN; Formal analysis and Visualization: RL, RN; Data interpretation: RL, RN, SH, SCF, HS; Supervision: SH, RN; Funding acquisition: SH, SCF, SR, MW, AEH, RN; Resources: SH, RN, SR, AEH, LE, MW; Writing -original draft: RL, RN; Writing – review & editing: RL, SR, LE, AEH, MW, SCF, HS, SH, RN. All authors have read and agreed to the published version of the manuscript.

## Acknowledgements

We thank Bettina Sonnenburg and Robert Günther for excellent technical assistance as well as Joshua Adjah for the mouse infections and Dr. Anne Winkler for the icons and sketches.

## Supplement Material

**Suppl.Fig. 1:**
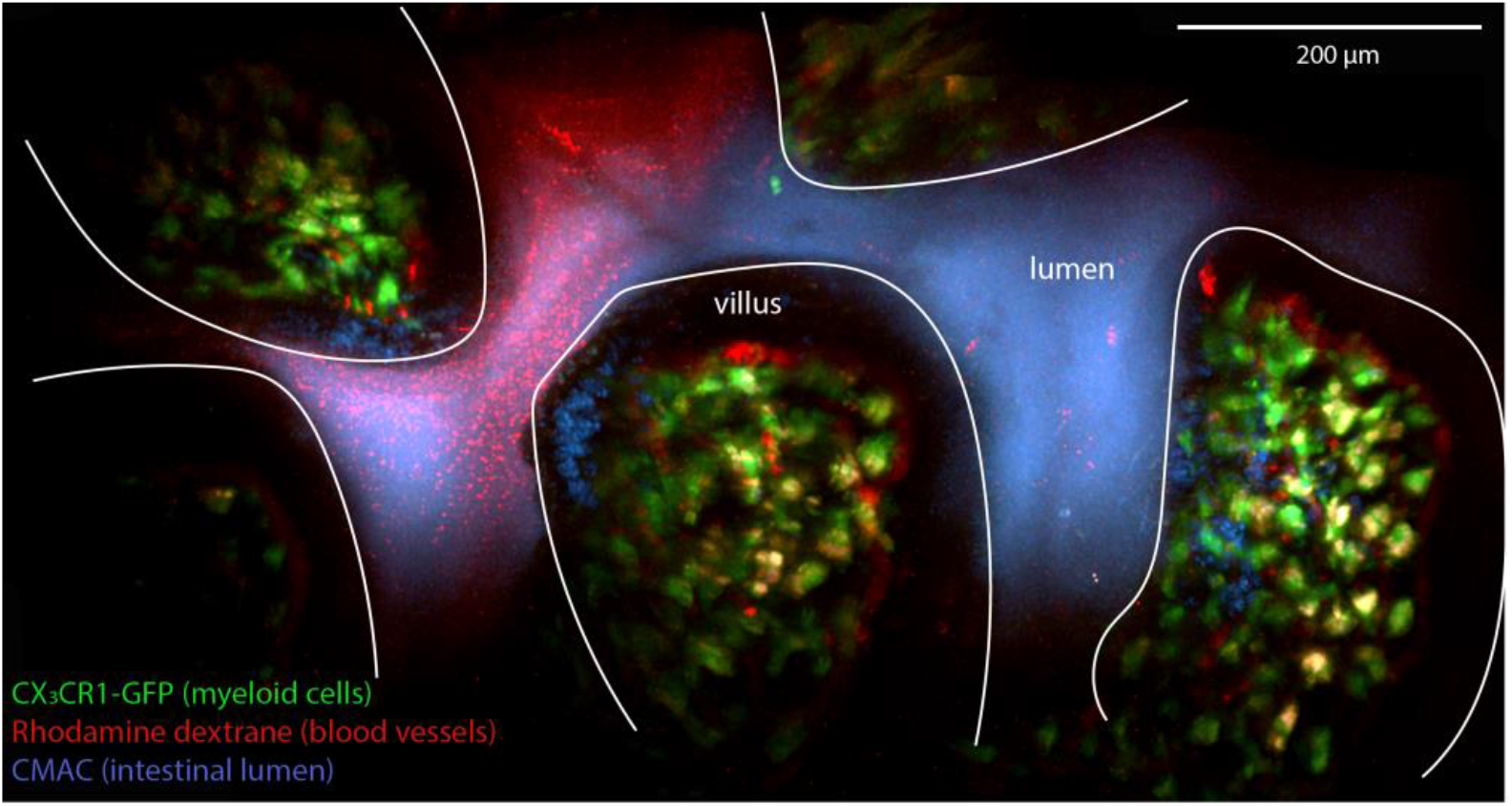
Two-photon microscopic image of the small intestine. Green: myeloid cells (CX3CR1-GFP), red: bood vessels (Rhodamine dextrane), blue: CMAC (intestinal lumen/ mucus).

**Suppl.Fig. 2:**
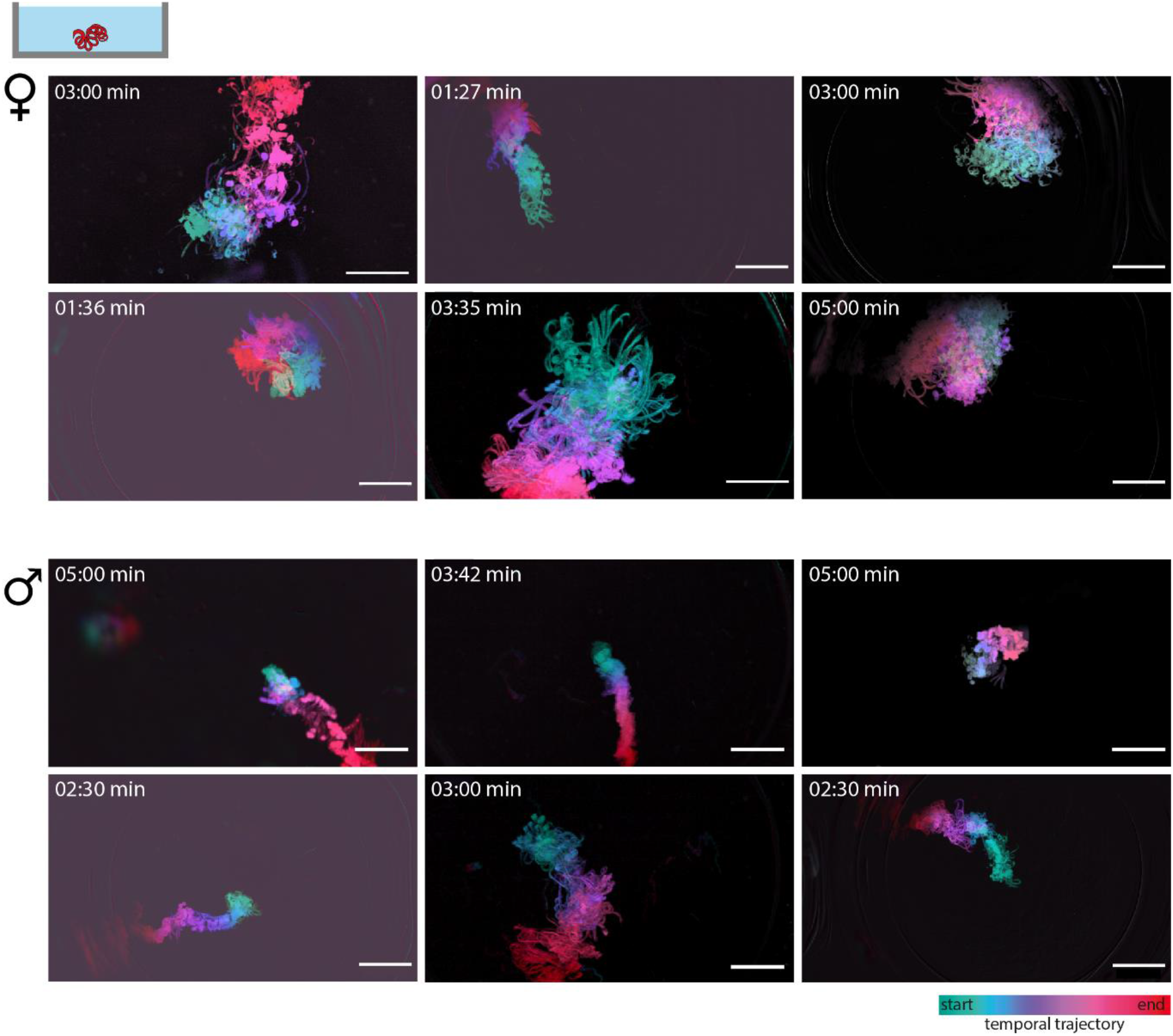
More exemplary temporal trajectories of *H.bakeri* motion under free-movement conditions depicting the variability of the movement behaviour. Top: femals, bottom: male. Scale bars indicate 3mm.

**Supple. Video 1: Living (right) vs. dead (left) *H. bakeri***. Mesoscopic movies at 0.67x magnification, played at 20x the normal speed, of worms in cell culture medium heated to 31°C. Shown dead nematodes were killed by low temperature (−20°C) and freshly defrosted and heated to 31°C for imaging. Time shown is hh:mm:ss; scale bar indicates 3 mm and applies for both movies.

**Supple. Video 2: Non-periodic motion pattern**. Exemplary microscopic movies at 4x magnification of a adult female (right) and a adult male (left) *H. bakeri*, videos are played at 10x the normal speed. Worms were in cell culture medium heated to 31°C. Time shown is hh:mm:ss; scale bars indicate 0.5 mm and applies for both movies.

**Supple. Video 3: Motion under unconfined conditions**. Exemplary mesoscopic movies at 0.67x magnification of an adult female (right) and an adult male (left) *H. bakeri* on a flat plate, played at 10x the normal speed. Worms were in cell culture medium heated to 31°C. Top: raw movies, bottom: processed data showing the evolving temporal trajectory. Time shown is hh:mm:ss, scale bars indicate 3 mm.

**Supple. Video 4: Motion in sex-mixed cohort under different conditions**. Mesoscopic movies (15 min) at 0.67x magnification, played at 20x the normal speed. Conditions: Left – worms in aqueous cell culture medium on a flat plate; Middle – worms in aqueous cell culture medium on 3D printed villi structure; Right – worms in viscous cell culture medium (2% alginate) on a flat plate, all media were heated to 31°C. Time shown is hh:mm:ss, scale bars indicate 3 mm.

**Supple. Video 5: Sex-mixed cohort 45 minutes in viscous, mucus-like medium**. Mesoscopic movie (45 min) at 0.67x magnification, played at 200x the normal speed. Worms in viscous cell culture medium thickened with 2% alginate on a flat plate, medium were heated to 31°C. Time shown is hh:mm:ss, scale bars indicate 3 mm.

**Supple. Video 6: Only short and sporadic interactions with 3D villi structure**. Mesoscopic movies at 0.67x magnification, played at 20x the normal speed. Worms in cell culture medium heated to 31°C and on 3D topography. The interactions with the structure are indicated with white arrows. Time shown is hh:mm:ss, scale bars indicate 3 mm.

